# Tandem split-GFP influenza A viruses for sensitive and accurate replication analyses

**DOI:** 10.1101/2025.07.27.667044

**Authors:** Himadri Nath, Ariel Arndt, Joshua T. Mann, Steven F. Baker

## Abstract

Reporter influenza A viruses (IAVs) are valuable tools for studying virus fitness, screening antivirals, and assessing host-virus interactions. However, the compact and segmented nature of the IAV genome presents major challenges for engineering genetically stable reporter viruses with minimal fitness defects. To address this, we developed a replication-competent IAV incorporating a tandem split-GFP reporter strategy. Specifically, we appended seven tandem repeats of the GFP11 peptide (GFP11×7) to the C-terminus of the PB2 or PA polymerase subunit. Viruses contained or lacked a self-cleaving 2A peptide between the viral and reporter genes to create polymerase fusion proteins or released GFP, respectively. These viruses are complemented in trans by cells expressing GFP1-10, allowing bright fluorescence with minimal disruption to viral function. The tandem GFP11 reporter viruses exhibit delayed replication kinetics, but are genetically stable over serial passages with a strong concordance between GFP signal and viral gene expression. We demonstrate their utility in high-throughput applications including fluorescence-based quantification of infection, neutralizing antibody titration, antiviral drug screening, and host factor identification, with results matching traditional assays but on an accelerated timeline. Furthermore, we engineered a battery of GFP1-10-expressing cell lines from multiple vertebrate species, illustrating the broad compatibility of this platform for comparative host studies. This system enables sensitive, scalable, and quantitative evaluation of influenza virus replication across diverse experimental contexts. The GFP11×7 reporter platform offers a versatile and robust tool for virology research and therapeutic screening, with potential for rapid adaptation to emerging IAV strains.

## INTRODUCTION

Influenza A virus (IAV) remains a persistent and significant global public health concern due to its high transmissibility, seasonal recurrence, and potential for pandemic emergence. Annual influenza epidemics result in approximately 1 billion infections globally, with an estimated 3 to 5 million cases of severe illness and 290,000 to 650,000 respiratory-related deaths each year^1^. In the United States alone, influenza has caused between 12,000 and 52,000 deaths annually over the past decade^2^. The risk escalates dramatically during pandemics; for instance, the 1918 H1N1 influenza pandemic is estimated to have caused at least 50 million deaths worldwide^3^. More recently, avian influenza viruses such as H5N1 and H7N9 have demonstrated zoonotic potential with alarming case fatality rates, reaching up to 60% for H5N1 and approximately 39% for H7N9^4,5^. Despite the availability of vaccines and antivirals, the ability of the virus to undergo antigenic drift and shift, coupled with its broad host range, underscores the urgent need for continued research into its biology, pathogenesis, and host interactions to inform better strategies for prevention and control.

IAV is an enveloped virus belonging to the *Orthomyxoviridae* family and possesses a segmented, negative-sense, single-stranded RNA genome. The genome comprises eight distinct RNA segments, each coding for one or more viral proteins, which are encapsidated by the nucleoprotein (NP) and bound at both termini by the heterotrimeric RNA-dependent RNA polymerase complex composed of the PB2, PB1, and PA subunits^6^. These viral ribonucleoprotein complexes (vRNPs) are essential for both transcription and replication of the viral genome. Infection begins via endocytosis when the virus receptor binding protein, hemagglutinin (HA), engages with sialic acid-containing glycoproteins on the host cell surface. Acidification of the endosome triggers HA-mediated fusion of the viral and endosomal membranes, leading to the release of vRNPs into the cytoplasm prior to their nuclear translocation^7^. vRNPs initiate mRNA synthesis using capped primers derived from host pre-mRNAs – a mechanism termed “cap-snatching”^7^. Following viral genome amplification, newly formed vRNPs are exported to the cytoplasm and directed toward the plasma membrane where viral assembly and budding take place. The selective packaging of all eight influenza A viral genome segments into progeny virions is a highly coordinated process thought to be governed by specific RNA-RNA and RNA-protein interactions. These interactions are mediated by cis-acting packaging signals that span the untranslated regions (UTRs) into the terminal coding sequences of each segment. While these signals are essential for efficient genome incorporation, the precise mechanisms underlying segment recognition and assembly remain only partially understood^8,9^.

Development of genetically encoded reporter IAVs has proven valuable for studying viral dynamics^10^. However, engineering such reporter viruses is challenging due to the compact (0.9 to 2.3 kb per segment) and highly constrained architecture of the IAV genome. To address this, we and others have demonstrated that segment-specific packaging signals can be functionally appended to the termini of modified gene segments, allowing the successful rescue of replication-competent viruses that harbor engineered sequences^11^. Two main types of reporters are commonly used: luminescent and fluorescent. While luminescent reporters offer high sensitivity for population-level assays, fluorescent reporters enable spatial and temporal tracking of infection at single-cell resolution. Insertion of full-length fluorescent proteins, such as enhanced GFP (eGFP), can disrupt packaging signals or impair viral protein function, resulting in attenuated replication, reduced fitness, or loss of reporter stability over serial passages^12–16^.

To circumvent size limitations for fluorescent influenza viruses, Avilov and colleagues developed a split-GFP virus that fused the C-terminal 16-amino-acid peptide of GFP (GFP11) onto the C-terminus of PB2^17^. This small tag is complemented *in trans* by the superfolder GFP1-10 molecule, stably expressed in host cells, allowing spontaneous and irreversible reconstitution of fluorescence with minimal perturbation to viral function^18,19^. We sought to strengthen the GFP signal in infected cells by inserting seven tandem repeats of GFP11 (GFP11×7) into the *PB2* or *PA* genes^18^. To preserve polymerase function, we introduced a self-cleaving 2A peptide from porcine teschovirus-1 (P2A) between the viral polymerase ORF and the GFP11×7 cassette. This ensures that the viral polymerase is released as a discrete, functional protein during translation, thereby minimizing potential interference with polymerase activity. As a result, the GFP11×7 virus retains replication kinetics, polymerase activity, and genetic stability. The simplicity and modularity of this strategy enables rapid generation of reporter viruses across multiple influenza strains, including newly emerging variants, without requiring strain-specific optimization. We demonstrate the utility of GFP11×7 viruses for antiviral screening, neutralization assays, and to identify species-specific host factors that contribute to replication success. This system enables quantitative assessment of host antiviral factors and visualization of infection dynamics at both population and single-cell levels. Collectively, GFP11×7 influenza viruses offer a robust and versatile platform to dissect critical steps of virus replication, investigate host-pathogen interactions, and evaluate species-specific barriers to infection.

## RESULTS

### Bicistronic polymerase-tandem GFP11 constructs have enhanced fluorescence and minimal replication defects

To generate reporter viruses that are accurate proxies for viral replication dynamics, the reporter genomic length, context, and protein size must be considered to preserve virus fitness. The split-GFP system allows a relatively short length (GFP11, 48 nt) to be incorporated into the viral genome, because engineered cells provide the bulkier remainder of (GFP1-10, 642 nt) of the transcript for complementation (**Fig 1A**). Moreover, the tandem repeats in GFP11×7 create a ∼10X brighter signal, while the length is still 60% that of full GFP (426 vs 714 nt) (**Fig 1B**)^18,20^. Both the PB2 and PA subunits of the viral polymerase have previously been shown to tolerate C-terminal fusions, and because these genes are expressed in low abundance compared to other common targets for reporters (eg. *NP, HA*, or *NS*), their linear expression profile can be assessed with high sensitivity to perturbations on viral gene expression^19,21^. We engineered several constructs to evaluate the impact on brightness and polymerase activity where GFP11 or GFP11×7 was fused to or released from PB2 or PA using a self-cleaving 2A peptide (P2A) (**Fig. 1C–D**)^22^. All constructs contained mutations at the wobble position in the codons spanning the terminal 109 or 50 nt of at the 3’ end of the ORF (PB2 or PA, respectively), because the packaging signals were duplicated after the reporter for subsequent virus rescue^23–25^. We first evaluated fluorescence signal intensity and localization in GFP1-10 overexpressing 293T cells (293T^GFP1-10^) transfected with plasmids expressing wildtype (WT) or modified *PB2* or *PA* genes. PB2 fused with a single copy of GFP11 (PB2-11) produced weak nuclear GFP signal, while the GFP11×7 fusion construct (PB2-11×7) yielded markedly stronger nuclear GFP signal, while the GFP11×7 fusion construct (PB2-11×7) yielded markedly stronger nuclear fluorescence, consistent with the nuclear subcellular localization of PB2 (**Fig. 1E**). In contrast, when the PB2-11×7 fusion was disrupted via a P2A peptide (PB2-2A-11×7), the signal was more diffuse and cytoplasmic, suggesting efficient separation of PB2 from the reporter tag. Similar GFP intensity and localization patterns were observed when Bicistronic *PA* produced GFP11×7 (PA-2A-11×7) in transfected cells.

**Figure 1.**
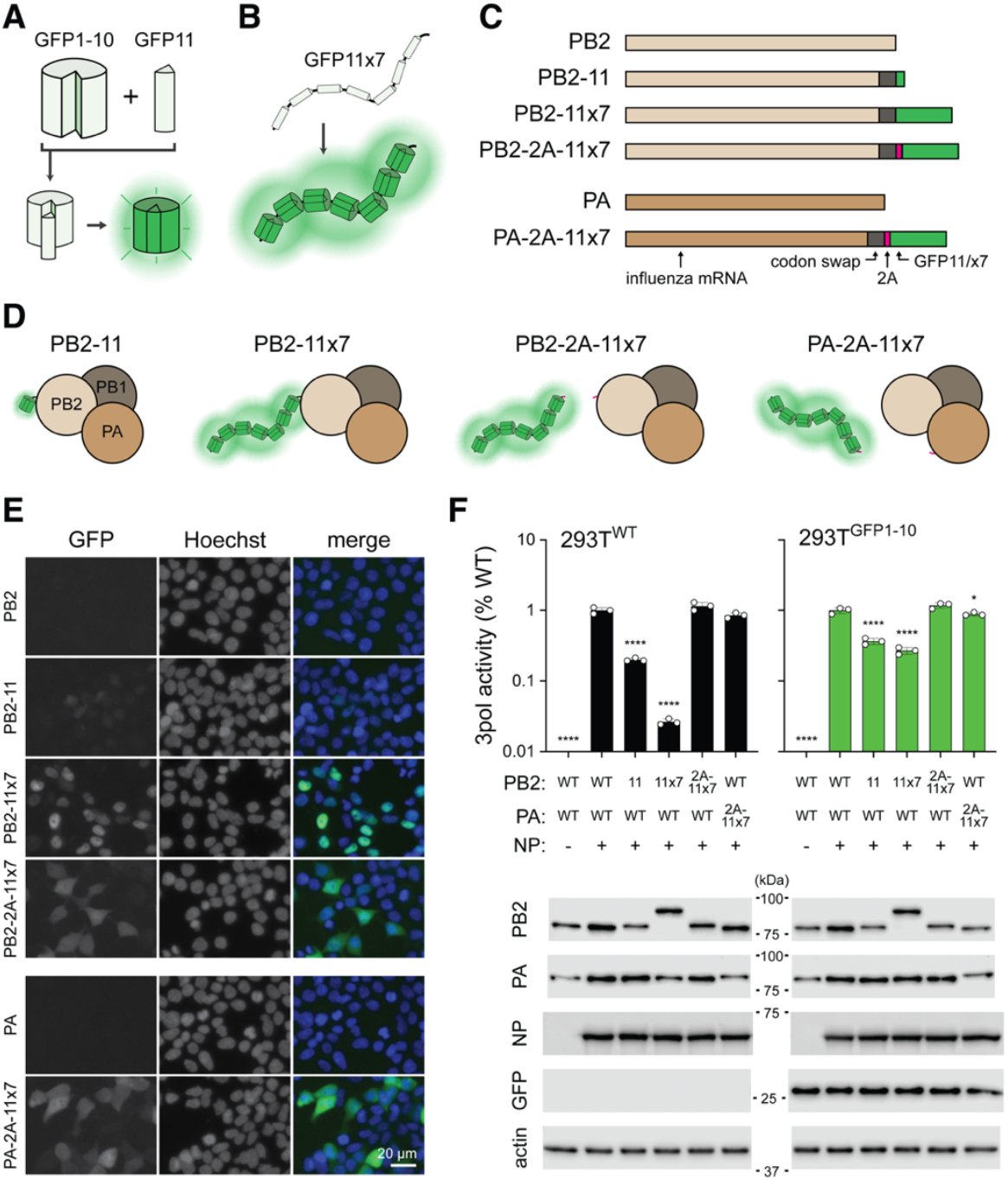
Virus polymerase genes with tandem repeats of GFP11. (A) Schematic of the self-assembling split-GFP system. (B) Strategy of tandem GFP11 repeats to increase fluorescence. (C) PB2 and PA reporter construct design with single or tandem GFP11, with or without a P2A self-cleaving peptide. Codon swap, silent mutations for packaging signal duplication. (D) Representations of different GFP11 reconstitution strategies with viral polymerase. Incorporation of P2A peptide ensures the separation of reporter from the polymerase complex. (E) Fluorescence microscopy of 293T^GFP1-10^ cells transfected with indicated constructs, nuclei stained with Hoechst. Scale bar, 20 µm. (F) Virus polymerase (3pol) activity assay measured in transfected 293T^WT/GFP1-10^ at 24 h with the indicated combinations of PB2, PA, and NP plasmids. All conditions additionally received PB1, firefly viral RNA reporter, and renilla pol II reporter. *Top*, normalized luminescence values shown with respect to WT condition. Representative experiment with *n* = 3 replicates. One-way ANOVA with Dunnett’s multiple comparisons test; ^*^, *P* < 0.05; ^****^, *P* < 0.0001. *Below*, 3pol activity lysates analyzed for protein expression by WB against the indicated proteins, kilodalton size (kDa) indicated.

To assess the functionality of these modified polymerase proteins, we performed minigenome assays in both 293T^WT^ and 293T^GFP1-10^ cells (**Fig 1F**). Cells were transfected with plasmids expressing WT or mutant polymerase genes in the presence or absence of NP. In both cell types, PB2 fusions with GFP11 or GFP11×7 led to significantly reduced polymerase activity. This reduction was more pronounced with GFP11×7 fusions, likely due to steric hindrance or altered protein stability^26^. However, incorporation of the 2A peptide (PB2-2A-11×7 or PA-2A-11×7) restored polymerase activity to near WT levels (**Fig. 1F**, top panel). Western blotting confirmed comparable expression levels of PA, PB2, and NP across different conditions and demonstrated PB2-fusions and efficient release of GFP11×7 from PB2 or PA in the 2A-separated versions (**Fig. 1F**, bottom panel). Together, these results validate our design strategy and demonstrate that tandem GFP11 tagging, in combination with 2A peptide separation, enables bright fluorescence while maintaining polymerase functionality.

### Characterization of tandem GFP11 viruses

To assess how the GFP11×7 reporter cassette affects viral replication, we generated lab-adapted influenza A/WSN/33 (H1N1; WSN) viruses carrying PB2-11×7, PB2-2A-11×7, or PA-2A-11×7 and evaluated their growth kinetics. Through plaque purification of PB2-11×7 virus rescue transfection, we recovered a virus with three tandem repeats of GFP11, where only the first of GFP11 is in frame. We refer to this virus as PB2-11x3*. Multicycle kinetics in MDCK cells show that all reporter viruses replicated efficiently but showed a delay and reduced peak titers compared to WT WSN virus (**Fig. 2A**). A major utility of GFP-expressing viruses lies within their potential analysis over single-cycle replication timescales, so we next evaluated viral gene expression in A549^GFP1-10^ cells over 8 h, representing one round of infection^27^. Gene expression of WT and tandem GFP11 viruses was measured by mRNA levels using RT-qPCR. All reporter viruses showed slightly reduced gene expression kinetics compared to WT, with a logarithmic increase in transcript abundance over the course of the 8 h infection (**Fig. 2B**). Importantly, replication of WT WSN was unaffected by the presence of GFP1-10 in A549^GFP1-10^ cells, as shown by similar multicycle kinetics in A549^WT^ and A549^GFP1-10^ cells, validating the substrate for our further biologic analyses (**Fig. 2C**). Next, we sought to evaluate whether GFP brightness could serve as a proxy for virus gene expression. We infected A549 cells with PA-2A-11×7 virus and measured both viral mRNA levels by RT-qPCR and the median fluorescence intensity (MFI) of GFP by flow cytometry over a single-cycle time course. MFI increased in parallel with viral mRNA levels between 2 and 8 h post-infection, indicating that the GFP11×7 reporter reflects viral gene expression (**Fig. 2D**). Notably, MFI intensity increases on a linear scale while mRNA abundance increases on a log_10_ scale.

**Figure 2.**
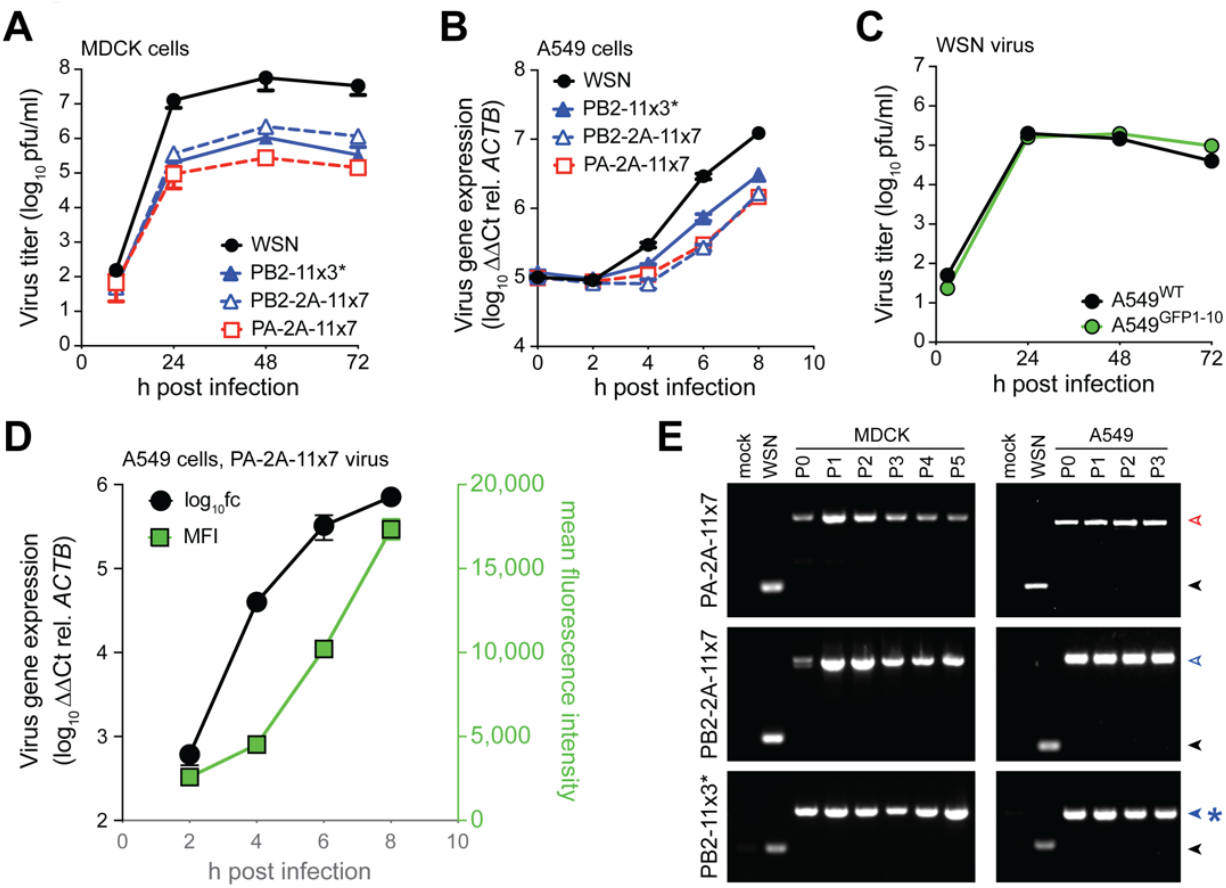
Growth kinetics and genetic stability of tandem GFP11 viruses. (A) Multicycle growth kinetics of WSN WT and tandem GFP11 reporter viruses in MDCK cells. Supernatants titrated by plaque assay. (B) Single cycle growth kinetics WT and tandem GFP11 reporter viruses analyzed by RT-qPCR for viral M mRNA accumulation in A549 cells. (C) Multicycle growth kinetics of WSN in A549^WT^ and A549^GFP1-10^ cells. (D) Dual analysis of viral gene expression over single cycle time course using RT-qPCR and flow cytometry. A549GFP^1-10^ cells infected with PA-2A-11×7 and analyzed at indicated times post-infection. (E) Genetic stability of reporter cassette analyzed by RT-PCR across multiple passages. MDCK or A549 cells were infected with indicated virus and blind passaged. RNA extracted from infected cells. Experiments were performed 2–4 independent times, each with technical replicates.

We next assessed the genetic stability of each reporter virus through multiple serial passages in MDCK and A549 cells. RT-PCR analysis of RNA from passage 0 to passage 5 confirmed stable retention of the GFP11×7 cassette in both PB2- and PA-tagged constructs. No loss or passaging-induced truncation of the reporter segment was detected (**Fig. 2E**), demonstrating the high genetic stability of these viruses after plaque purification. Due to the fact that PA-2A-11×7 virus was easy to rescue (3/3 positive wells at P1 from rescue) compared to PB2-2A-11×7 (1/3 positive wells at P2 from rescue), and that PB2-11x3* was artificially selected by plaque isolation – most of the rescued virus that induced fluorescence did not plaque – we continued with PA-2A-11×7 for further experiments. Altogether, these results demonstrate that tandem GFP11 reporter viruses retain replication competence, are genetically stable, and the fluorescence signal can be used as a proxy for viral gene expression.

### High-throughput-compatible infection quantitation to evaluate antivirals

We first optimized a plate reader-based assay of GFP11×7 virus infection and determined the linear range of detectable signal. MDCK-TMPRSS2^GFP1-10^ cells^28^ in 96 well plates were infected with serial dilutions of PA-2A-11×7 virus, and fluorescence was measured in live infected cells 24 h post-infection. The GFP signal peaked when cells were infected with ∼1,280 PFU (MOI = 0.043), representing a 10-fold increase in signal intensity over background. Moreover, we observe a sigmoidal dose-response relationship with a calculated EC_50_ of 284 PFU (**Fig. 3B**). Next, we assessed the application of this system for neutralization assays by testing the ability of anti-influenza antibodies to block infection. We pre-incubated 500 PFU (MOI, 0.017) of PA-2A-11×7 with serial dilutions of antibodies generated against matched WSN (NR-3104) or mismatched influenza B/Brisbane/60/2008 (NR-58872). As expected, NR-3104 exhibited potent neutralization activity, with complete inhibition through a 1:800 dilution of serum and a calculated NT_50_ of 1:1,018. We benchmarked our GFP-based assay with classic CPE-based neutralization assays using WT virus, which after four days yielded a neutralization titer for NR-3104 of 1:800. In contrast, the influenza B specific antibody NR-58872 showed no measurable GFP-based (**Fig. 3C**) or CPE-based neutralizing activity. These results validate the ability of GFP11×7 virus to accurately detect neutralizing antibody responses in vitro using a fast and sensitive fluorescence-based assay.

**Figure 3.**
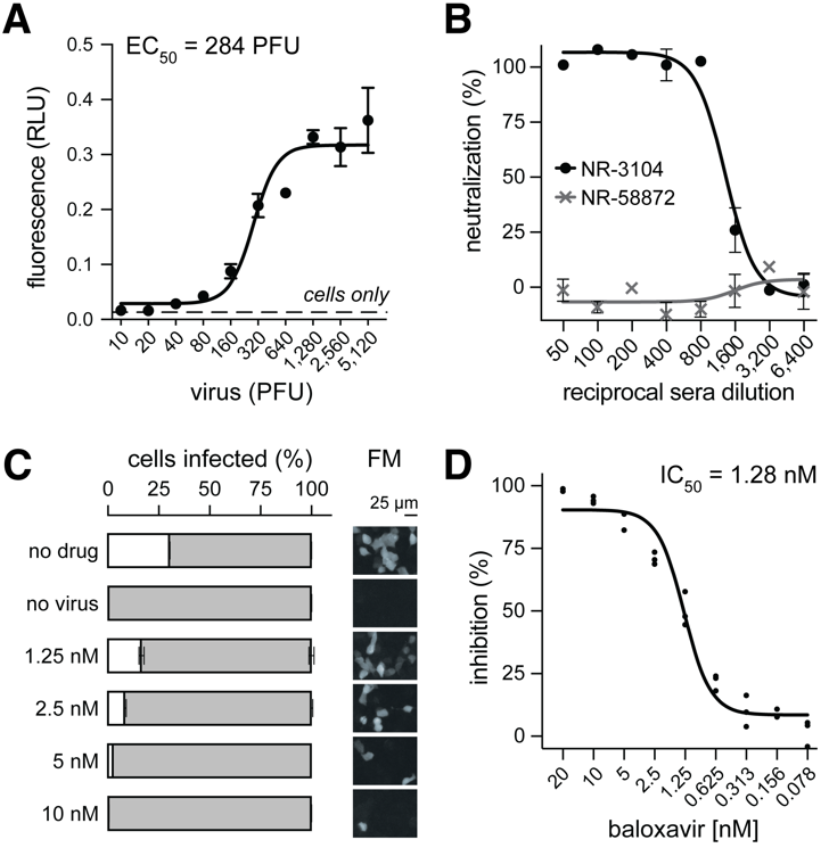
Platforms to identify virus neutralizing antibodies and antiviral compounds. (A) Fluorescence-based plate reader assay of MDCK-TMPRSS2^GFP1-10^ cells infected with PA-2A-11×7 virus for 24 h. (B) Microneutralization assay as in (A), where 500 PFU PA-2A-11×7 virus was pre-incubated with dilutions of influenza A-specific serum (NR-3104) or influenza B-specific monoclonal antibody (NR-58872). GFP readings taken 24 h post-infection. (C-D) Antiviral drug sensitivity assay. Baloxavir marboxil treatment in infected cells for 24 h. C) Flow cytometric percentage of infection correlates with reduction in GFP cells detected by fluorescence microscopy (FM). Scale bar, 25 µm. D) Dose-response inhibition of baloxavir over *n* = 3 independent experiments. Infection percentage quantified by flow cytometry.

To demonstrate the utility of the reporter virus for antiviral drug screening, we evaluated its sensitivity to baloxavir marboxil, a cap-dependent endonuclease inhibitor used clinically for influenza treatment^29^. A549^GFP1-10^ cells were infected with PA-2A-11×7 virus in the presence of increasing concentrations of baloxavir and incubated for 24 h. Flow cytometry analysis revealed a dose-dependent reduction in GFP-positive cells (**Fig.3E**), with complete inhibition at a concentration of 10 nM baloxavir. Fluorescence microscopy confirmed this pattern, with a marked reduction in GFP-expressing cells at baloxavir concentrations above 2.5 nM (**Fig. 3E**, right panel). Dose-response assays across multiple experiments revealed highly reproducible values, with an IC_50_ value of 1.28 nM (**Fig. 3F**), which is consistent with previously reported values of 1.3 to 1.6 nM^30^.

### Analysis of host factors using GFP11×7 influenza virus

Influenza A viruses have a broad host tropism, but species-specific differences of pro or antiviral factors play a major role in determining the replicative ability across hosts. Because the split-GFP platform requires cells that constitutively express GFP1-10, we engineered a menagerie of cell lines representing the diverse host phylogeny of influenza and influenza-like virus hosts (**Fig. 4A**). This panel of GFP1-10-expressing vertebrate cells includes human, primate, bovine, canine, porcine, and avian cell lines. Upon infection with PA-2A-11×7, all cell lines displayed strong GFP fluorescence, confirming that the reporter virus is compatible with multiple host backgrounds and that the GFP signal faithfully tracks viral replication across species. This establishes a reliable platform for monitoring influenza virus infection dynamics across diverse cell types.

**Figure 4.**
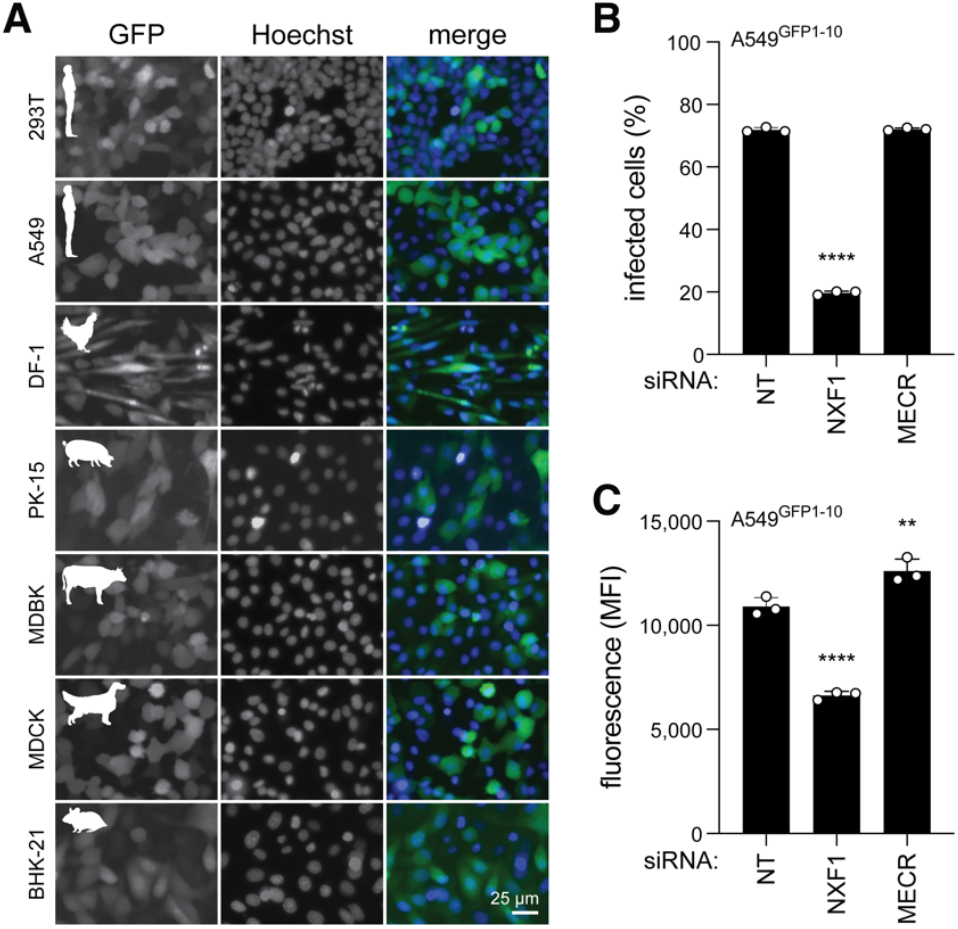
Screening compatibility of host gene perturbations using influenza GFP11×7. (A) GFP1-10-expressing cell lines derived from human, avian, porcine, bovine, canine and rodent species show GFP signal when infected with PA-2A-11×7 reporter virus,. Various MOIs of 0.05 – 0.005, 24 h infection. Scale bar, 25 µm. (B-C) A549^GFP1-10^ cells transfected with siRNA for 48 h prior to 6 h infection with PA-2A-11×7. Analysis of percentage and median fluorescence intensity of GFP signal by flow cytometry. B) Percentage of infected cells. C) Intensity of reporter as proxy for virus gene expression. Representative experiment with *n* = 3 replicates. One-way ANOVA with Dunnett’s multiple comparisons test r; ^**^, *P* < 0.01; ^****^, *P* < 0.0001.

To evaluate the utility of the GFP11×7 reporter virus for screening perturbations of specific host factors involved in influenza virus replication, we performed siRNA-mediated knockdown of genes with known pro- or antiviral functions. Specifically, we targeted *NXF1*, a pro-viral host factor^31,32^, and *MECR*, an antiviral gene^33^ in human A549^GFP1-10^. At 6 h post-infection, fluorescence intensity was quantified to assess viral replication.

Knockdown of *NXF1* led to a marked reduction in percentage of cells infected and MFI, consistent with impaired viral replication (**Fig. 4B**). In contrast, silencing of *MECR* resulted in no change in percentage of infected cells, but a significant increase in MFI compared to control, reflecting enhanced virus replication due to loss of an antiviral barrier. These results demonstrate that the reporter virus can sensitively detect the impact of host gene modulation within a single cycle of infection. The rapid and quantitative readout supports its application in functional genetic screens, including siRNA and CRISPR-based approaches, to identify both pro- and antiviral host factors.

## DISCUSSION

Mitigating the public threat posed by influenza viruses requires a diverse portfolio of approaches, ranging from screening existing FDA-approved drugs for off-label antiviral activity to advancing our understanding of host– virus interactions. Furthermore, a robust understanding of virus-host interactions and infection dynamics is essential to develop novel therapeutic strategies. In this study, we present an optimized reporter system for IAV that addresses long-standing limitations in reporter virus design. By leveraging a split-GFP strategy and incorporating tandem repeats of GFP11 with a self-cleaving 2A peptide, we generated a replication-competent reporter virus that maintains near-WT polymerase activity, with a modest reduction in growth kinetics, while maintaining strong genetic stability. This system enables sensitive detection and visualization of viral infection in living cells at the single cell level.

Our split GFP approach requires less genetic space, and packaging signal duplication and codon swapping limits viral elimination of the reporter cassette. Interestingly, the PB2-11x3* virus was generated through random mutation and selection. Sequencing revealed that instead of seven tandem repeats of GFP11, PB2-11x3* virus instead has PB2 fused to one GFP11 immediately followed by an additional two GFP11 repeats in the +1 ORF. Despite this aberrant genome architecture, GFP1-10 cells infected with PB2-11x3* exhibit a bright, nuclear signal (data not shown). We speculate that this variant was selected for because the majority of PB2-11×7 virus rescue supernatant induce bright, nuclear GFP expression but was unable to form plaques. PB2-11x3* virus was serially plaque purified three times, selecting only GFP+ clones. Because this virus is much brighter than PB2-11 yet retains nuclear GFP signal, we hypothesize that deletion-containing viral genome-encoded proteins are being synthesized that link the N-terminus of PB2 with the out of frame two tandem repeats of GFP11^34^. We also rescued PA-2A-11×7 viruses in other backbones of influenza A – CA07 and PR8 (data not shown) – suggesting this platform can be rapidly deployed to perform quantitative analyses of newly emerging strains.

We demonstrated the potential of this reporter virus for high-throughput applications. The strong GFP signal allows sensitive quantification of infection using a plate reader or flow cytometry to evaluate neutralizing antibodies or antiviral compounds. The dose-dependent inhibition of fluorescence signal by NR-3104 or baloxavir was consistent with prior assays that require 4-5 days rather than the 24 hours needed here. Beyond antiviral screening, this reporter virus shows significant potential for identifying host factors involved in viral replication. siRNA-mediated knockdown revealed pro-viral and antiviral gene functions within 6 hours. This rapid and sensitive detection underscores the utility of the GFP11×7 virus as a platform to dissect host-virus interactions or to perform functional genomics in a variety of host species.

The GFP11×7 virus offers several advantages over traditional influenza reporters. First, it provides a real-time, live-cell readout without the need for fixation, chemical labeling, or a secondary reactant. Second, it is compatible with multiple detection formats; we show here microscopy, flow cytometry, and plate readers, but this should also be amenable to similar technologies like confocal, quantitative, or high-content live cell imaging. Third, it maintains robust replication and genetic stability. Fourth, it can rapidly be applied to other influenza strains due to the ease of reverse genetics and rescue. Fifth, the streamlined 6 or 24 hour assays offer fast timelines to evaluate host factors or neutralizing antibody and antiviral compounds, respectively. Lacking, however, is a suitable GFP1-10 animal model to study live imaging *in* or *ex vivo*, although a transgenic chicken may soon be reported (H. Sang, Roslin Institute). Developing live animal assays with GFP11×7 viruses would be beacon to illuminate basic organismal influenza virology.

In summary, we present a bright influenza A split-GFP virus that recapitulates WT-like replication dynamics over single-cycle timescales. This tool enables dynamic tracking of infection, quantitative antiviral or neutralizing antibody screening, and host-pathogen interaction studies across diverse species. Its flexibility and scalability make it a powerful platform for both basic research and translational virology efforts directed toward pandemic preparedness.

## ACKNOWLEDGEMENTS

We thank members of the Baker lab for vibrant discussion and critical reading of the manuscript. We thank A. Mehle (University of Wisconsin-Madison), for sharing influenza reverse genetics plasmids and anti-PB2 antibody. Antibodies (NR-3133, NR-3104, and NR-57882) were kindly provided by Bei Resources, NIH. GFP1-10 plasmid kindly provided by Addgene (70219). This work was supported by the American Lung Association (ERPALA2023).

## MATERIALS AND METHODS

### Cell culture

A549, 293T, MDCK, MDCK-TMPRSS2, MDBK, BHK21, DF-1, and PK-15 cells were maintained in Dulbecco’s Modified Eagle Medium (DMEM; Gibco) supplemented with 10% fetal bovine serum (FBS), and incubated at 37°C with 5% CO_2_, except for DF-1 cells, which were maintained at 39°C. Cells were routinely evaluated for mycoplasma (Lonza). For viral infections, virus growth medium (VGM) was used, consisting of DMEM supplemented with 0.2% bovine serum albumin (BSA), 25 mM HEPES, 1x penicillin-streptomycin, and TPCK-treated trypsin (0.25-2 µg/mL; Sigma). Stable cell lines expressing GFP1-10 were generated via retroviral transduction. Transduced cells were selected using appropriate antibiotics (puromycin or neomycin).

### Plasmid construction

Reporter rescue plasmids encoding PA or PB2 fused with tandem GFP11 segments were constructed using Gibson assembly (New England Biolabs) or inverse PCR, following prior principles^17,25^. Constructs included either a single GFP11 peptide or seven tandem repeats (GFP11×7), fused in-frame to the C-terminus of PA or PB2 open reading frames (ORFs). In some constructs, a porcine teschovirus-1 2A (P2A) self-cleaving peptide was inserted between the viral ORF and GFP11×7 to ensure post-translational separation of the two proteins. To preserve segment packaging, packaging signals (5’ terminal [genomic sense] 109 nt of PB2 or 50 nt of PA) were duplicated between the stop codon and native 5′ UTR. Silent mutations were introduced into the packaging signals that overlap with PB2 or PA coding sequences. pQCXIP GFP1-10 was generated using Gibson assembly with pcDNA3.1 GFP1-10 and pQCXIP/N.

### Virus rescue and titration

Influenza A viruses were generated by reverse genetics: A/WSN/1933 (H1N1; WSN), A/Puerto Rico/8/1934 (H1N1; PR8) A/California/07/2009 (H1N1; CA07). 293T cells were transfected with eight pBD or pHW2000 viral genome plasmids (PB2, PB1, PA, HA, NP, NA, M, NS) from various influenza A viruses and supporting expression plasmids (p3X-1T, pCAGGS NP, TMPRSS2) using jetPRIME (Polyplus). MDCK TMPRSS2 cells were overlayed one day post-transfection to facilitate viral amplification^35^. Reporter viruses were generated by replacing WT PB2 or PA segments with respective reporter constructs. Rescued viruses were plaque purified on MDCK cells, sequence-confirmed by Oxford Nanopore (Plasmidsaurus), and titrated by plaque assay on MDCK cells.

### Polymerase activity assay

293T WT or GFP1-10 cells were seeded in 24-well plates and transfected in triplicate with plasmids encoding PB1, WT or GFP11 versions of PB2 or PA, and NP or empty vector using TransIT-X2 (Mirus). A firefly luciferase reporter plasmid (pHH21-vNA-Luc), flanked by influenza virus UTRs, was co-transfected to assess viral polymerase activity^36^. A constitutively expressed renilla luciferase plasmid served as a normalization control (Promega). 24 h post-transfection, cells were lysed, and firefly and renilla luciferase activity were measured. Lysates were also analyzed by western blot to verify expression and cleavage of polymerase and reporter proteins using the following antibodies: rabbit anti-PB2^37^; rabbit anti-PA, GeneTex GTX125933; goat anti-NP, BEI NR-3133; rat anti-GFP, Chromotek 3H9; rabbit anti-ß-actin, GeneTex GTX637675.

### Microscopy

GFP fluorescence driven by reporter constructs was evaluated by transfecting 293T^GFP1-10^ cells. At 24 h post-transfection, cultures were incubated with Hoechst 33258 (Invitrogen) diluted in culture medium for 30 minutes following manufacturer instructions. After incubation, the Hoechst-containing medium was replaced with 1X DPBS. For infected cell imaging, at 24 h post-infection, Hoechst stain was added as above. Images were captured with a 20X objective on an EVOS M5000 (Thermo Fisher)**Multicycle growth kinetics** Viral replication kinetics were evaluated in A549 or MDCK cells by infecting at a multiplicity of infection (MOI) of 0.01. Supernatants were collected at designated time points post-infection and stored at -80°C until titration. Viral titers were determined by plaque assay on MDCK cells^38^.

### Single-cycle kinetics by RT-qPCR and flow cytometry

For single-cycle replication analysis, cells were infected at an MOI of 0.2. Triplicate infected cells were stored in TRIzol every 2 h from 0 to 8 h post-infection. Total RNA was extracted prior to RT-qPCR using primers specific for viral M mRNA^39^ (Luna, NEB). Human β-actin mRNA was used for normalization via the ΔΔCt method. Primer sequences are as follows: Flu_qPCR_FOR, ATGAGYCTTYTAACCGAGGTCGAAACG; Flu_qPCR_REV, TGGACAAANCGTCTACGCTGCAG; ActinB_FOR, CTGGAACGGTGAAGGTGACA; ActinB_REV, AAGGGACTTCCTGTAACAATGCA. For fluorescence quantification, parallel samples were fixed and analyzed by flow cytometry to measure GFP median fluorescence intensity (MFI).

### Reporter virus stability across passages

Reporter viruses were used to initially infect at an MOI of 0.1 (A549) or 0.01 (MDCK) and subsequently blind passaged serially 2-4 times. Total RNA was extracted from infected cells at each passage, and RT-PCR (Superscript III, Invitrogen; OneTaq Hot Start DNA Polymerase, NEB) was performed to amplify the reporter-encoding region and confirm retention of the GFP11×7 cassette. The primer sequences are as follows: PA int FOR,GTTCAGGCTCTTAGGGACAAC; Bm-PA-2233R, ATATCGTCTCGTATTAGTAGAAACAAGGTACTT; PB2 int FOR,CAGGAGATATGGACCAGCATTAAG; Ba-PB2-2341R, ATATGGTCTCGTATTAGTAGAAACAAGGTCGTTT^40^.

### Virus neutralization assay

Reporter virus (500 PFU, 0.017 MOI) was pre-incubated with serial dilutions of serum (NR-3104, specific to WSN) or monoclonal antibody (NR-58872, specific to influenza B HA) for 1 h at 37°C in VGM. Virus-antibody mixtures were then added to 96-well black-bottom plates seeded with MDCK-TMPRSS2^GFP1-10^ cells. After 24 h, wells were washed with 1X DPBS and GFP fluorescence was measured using a plate reader (Fluoroskan-FL, Thermo Fisher). Neutralization calculations performed using the following formula:

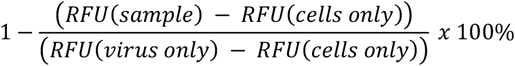

### Drug inhibition assay

A549^GFP1-10^ cells were incubated with the reporter virus (MOI, 0.1) and increasing concentrations of baloxavir marboxil (Selleckchem) for 24 h. Afterwards, cells were trypsinized, fixed with 4% paraformaldehyde, and analyzed by flow cytometry (BD FACSCelesta). The percentage of GFP-positive cells was quantified to assess infection inhibition (FlowJo). Representative wells were also imaged via fluorescence microscopy.

### siRNA-mediated knockdown of host factors

A549^GFP1-10^ cells were transfected with either non-targeting or gene-specific siRNAs targeting *NXF1* and *MECR* (SMARTpool, Dharmacon) using TransIT-X2. At 48 h post-transfection, cells were washed with VGM and infected with the PA-2A-11×7 virus (MOI, 0.2). 6 h post-infection, cells were processed for flow cytometry as above to measure percent infected MFI of GFP as a readout of viral replication (FlowJo). Percent inhibition performed using the following formula:

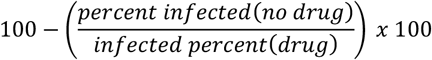

### Statistics

Experiments were performed 2-4 independent times each containing technical replicates. Data presented are either a representative experiment or the average of multiple biologic replicates (grand mean). Statistics performed using Prism (GraphPad).

